# Mitotic kinase regulation of DNA replication forks

**DOI:** 10.64898/2026.05.16.725582

**Authors:** Sudikchya Shrestha, Natasia Paukovich, Samuel R. Greenfield, Andrea MacFadden, Shania N. Smith, Shaun Bevers, Allison W. McClure

## Abstract

While DNA replication forks initiate in S phase, they do not necessarily complete and terminate prior to cell entry into mitosis. How mitotic proteins regulate leftover replication forks is not well-understood. Using reconstituted DNA replication forks with purified proteins, we show that the budding yeast mitotic kinases Clb2-CDK (M-CDK) and Cdc5 (Plk1 homolog) phosphorylate and regulate several replication elongation proteins. Mrc1 phosphorylation by both kinases results in slower replication, and Polα phosphorylation by M-CDK results in less lagging strand initiation. We further show that a phospho-resistant mutant of Polα bypasses M-CDK inhibition of Polα activity in reconstituted replication reactions. Yeast cells expressing the phospho-resistant mutant exhibit faster cell cycle progression revealing a potential negative feedback mechanism between DNA replication forks and mitotic progression.

## Introduction

The bulk of DNA replication occurs in S phase and is temporally separated from mitosis when chromosomes are segregated into progeny cells. However, there is no known mechanism that ensures replication forks finish replicating the entire genome prior to the cells moving on to G2 phase and mitosis. Evidence from human and budding yeast shows that cells can enter mitosis with unreplicated DNA that is then synthesized in mitosis^1–3^ (which has been called MiDAS for <u>mi</u>totic <u>D</u>N<u>As</u>ynthesis). If unreplicated DNA is not resolved in mitosis, anaphase bridges form during chromosome segregation, causing genome instability and frequently leading to aneuploidy or mitotic failure, hallmarks of cancer^4–6^.

DNA replication is initiated at origins of replication where MCM hexamers are loaded in the G1 phase of the cell cycle (reviewed in ^7^). Upon entry into S phase, these MCM hexamers are converted into active CMG helicases and assemble with the replicative polymerases and accessory factors to form the replisome. The replisome unwinds parental DNA and synthesizes nascent strands on the exposed ssDNA template. Replication is terminated when two replication forks converge triggering replisome disassembly^8^.

If a replication fork does not finish replicating and terminate within S phase, what happens when cells move on to G2 and mitosis? These “leftover” forks may not finish for many reasons including blocks in the DNA template, e.g. protein roadblocks or crosslinked parental DNA, or because there are not enough replication forks to replicate a long stretch of DNA within the S phase timeframe^9^. Classic studies showed that fusing S phase and mitotic cells or prematurely inducing mitosis triggers dramatic DNA fragmentation at replication forks^10^. These findings established the long-held view that the mitotic environment is inhospitable to S phase replication forks. However, this is hard to reconcile with the reports of cells entering mitosis with unreplicated DNA and subsequently completing synthesis.

Studies in *Xenopus* egg extracts and in mammalian cells have identified a mitotic pathway to remove CMG from these unfinished replication forks^7–10^. Phosphorylation by mitotic cyclin-dependent kinase (M-CDK) of the E3 ubiquitin ligase TRAIP is necessary for CMG disassembly in mitosis in metazoans, but artificial phosphorylation of TRAIP is not sufficient to trigger CMG disassembly in S phase^11^. Therefore, other changes that occur only in mitosis must be required for CMG removal and replication fork processing to allow or promote MiDAS.

Entry into mitosis is characterized by the rise of M-CDK and Polo-like kinase activity. In budding yeast, M-CDK is composed of the kinase Cdc28, the substrate adaptor Cks1, and either Clb2 or Clb1 cyclin. The polo-like kinase homolog in budding yeast is Cdc5. Each of these kinases has hundreds of targets^12–17^, so parsing individual phosphorylation events has historically been difficult. In this study, we leveraged the biochemical reconstitution of DNA replication to understand how replication forks are regulated by M-CDK and Cdc5. We identified two functional targets of the mitotic kinases: Mrc1 and Polα, leading to a reduction in replication elongation rate and lagging strand initiation, respectively. Using a phospho-resistant mutant of Polα, we also identified Polα phosphorylation as a mechanism regulating mitotic progression.

## Results

### Mitotic kinases slow replication restart following reversible pause

To understand how mitotic kinases affect replication forks that have not finished when cells enter mitosis, we paused reconstituted replication forks, added purified mitotic kinases, and then resumed replication. Replication proteins were expressed from budding yeast strains modified with the β-estradiol-sensitive hybrid transcription factor *GAL4*^*DBD*^*-hER-VP16*^*AD*18^ (see supplemental methods) and then purified. Replication forks were then biochemically reconstituted^19,20^: replication was initiated on a plasmid template containing the ARS1 budding yeast origin in the presence of a pulse of radiolabeled dCTP allowing selective visualization of newly synthesized DNA. Okazaki fragment nuclease and ligase were omitted to allow for visualization of leading and lagging strands separately.

In one pausing strategy, topoisomerases were omitted from the replication reaction^21^, which prevented replication from proceeding further than ∼1 kb on the 10 kb plasmid template (Fig 1A, lane 1). Addition of purified Top1 readily restarted replication synchronously (Fig 1A, lanes 4-5). However, topoisomerase omission only paused forks in reactions containing low salt (100 mM potassium glutamate).

**Figure 1.**
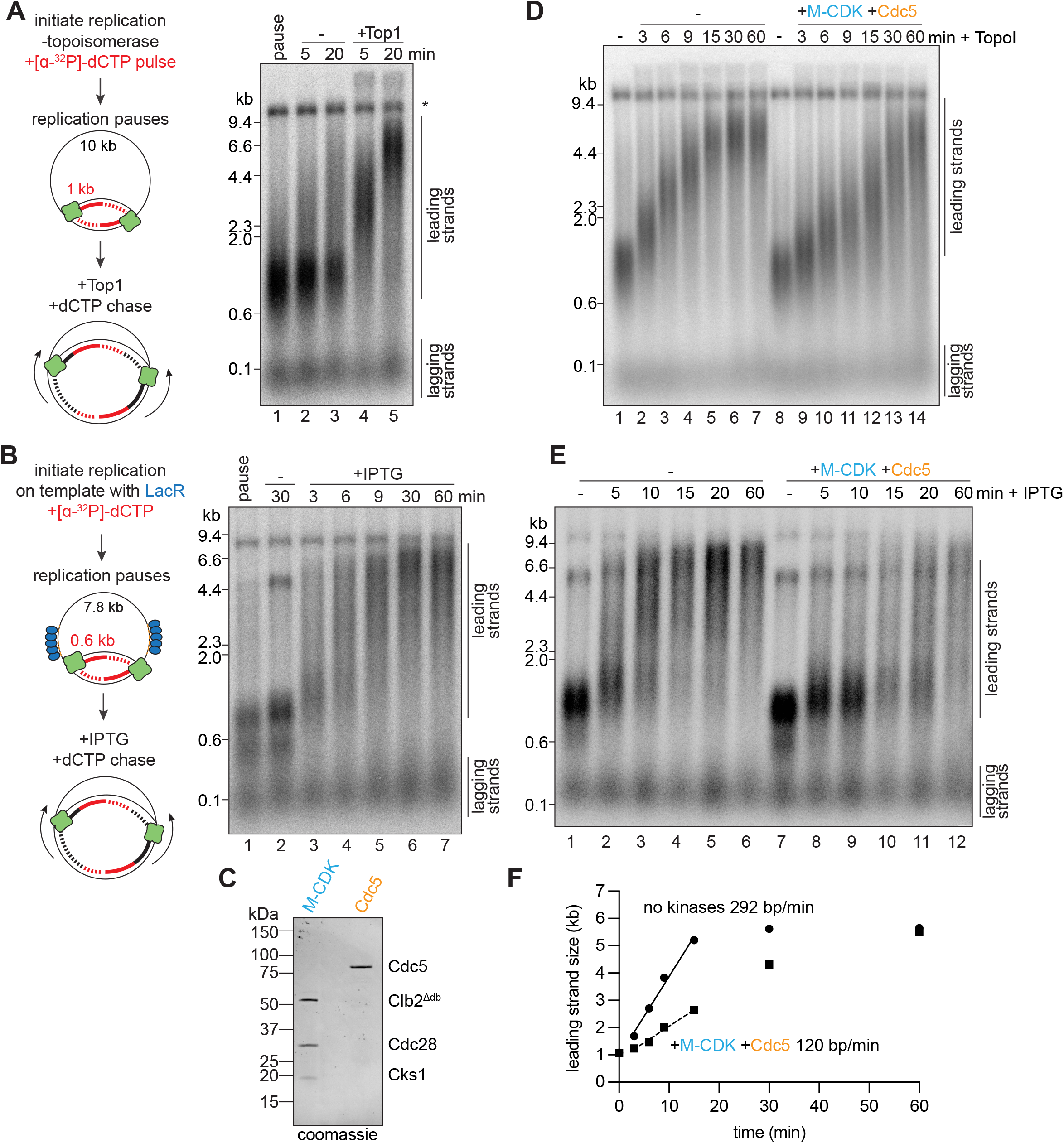
Mitotic kinases slow DNA replication restart following reversible pause. A) DNA replication was reconstituted with purified proteins on ARS1 plasmid (pJY22) in the absence of topoisomerases leading to a pause around 1 kb. Replication was resumed upon the addition of Top1 with excess unlabeled dCTP. * indicates replication-independent template labeling. B) Template with artificial ARS flanked by lacO arrays (pAWM197) was pre-incubated with LacR prior to initiation of DNA replication. Replication was resumed upon addition of IPTG. C) Purified Cdc5 and M-CDK were separated on SDS-PAGE and coomassie stained. D) M-CDK and Cdc5 were added to paused replication forks (as in A) and then replication was resumed with TopoI. E) M-CDK and Cdc5 were added to paused replication forks (as in B) and then replication was resumed with IPTG. F) quantification of bulk leading strand sizes from D.

The second strategy for pausing reconstituted replication forks made use of protein-DNA barriers^22,23^. Replication was initiated on a 7.8 kb DNA plasmid containing 16X and 19X repeats of the *lacO* sequence bound by LacR at a distance of ∼0.6 kb on either side of the replication origin sequence (Fig 1B). Replication was then restarted by the addition of IPTG, which disrupts LacR binding (Fig 1B). Restart was less synchronous than the topoisomerase omission strategy, but the reactions could be performed in higher salt conditions (250 mM potassium glutamate).

To interrogate the effect of mitotic kinase regulation on paused replication forks, we purified Cdc5 and M-CDK from budding yeast synchronized in prometaphase with nocodazole (Fig 1C). M-CDK was purified as the Cdc28-Cks1-Clb2 trimer with the Clb2 cyclin degron box deleted to increase expression. We incubated paused replication forks with Cdc5 and M-CDK, then restarted replication. Replication reactions were pulse labeled, so only existing replication fork DNA was visualized. Following mitotic kinase incubation, paused forks formed by topoisomerase omission or LacR block readily restarted and achieved full-length products (Fig 1D, E). However, the elongation rate in reactions with mitotic kinases was slower (120 bp/min) compared to controls incubated with buffer (292 bp/min) (Fig 1F).

### Mitotic kinases phosphorylate Mrc1 to slow replication

To identify candidate protein(s) at the replication fork that could be targeted by the mitotic kinases to slow replication, we incubated Cdc5 or M-CDK individually with the known elongation regulators Mrc1, Csm3, and Tof1^7,20,24^. Cdc5 induced slower migration in an SDS-PAGE gel of Mrc1, Csm3, and Tof1 consistent with phosphorylation (Fig 2A). M-CDK induced slower migration of Mrc1 but not Csm3 or Tof1 (Fig 2A). To directly test whether phosphorylation of these targets was responsible for replication slowing, we pre-incubated them with M-CDK and Cdc5 prior to addition to the replication reaction. Mrc1 phosphorylated by either Cdc5 or M-CDK slowed replication (Fig 2B,C), but Csm3 and Tof1 pre-incubation with Cdc5 did not slow replication (Fig 2D).

**Figure 2.**
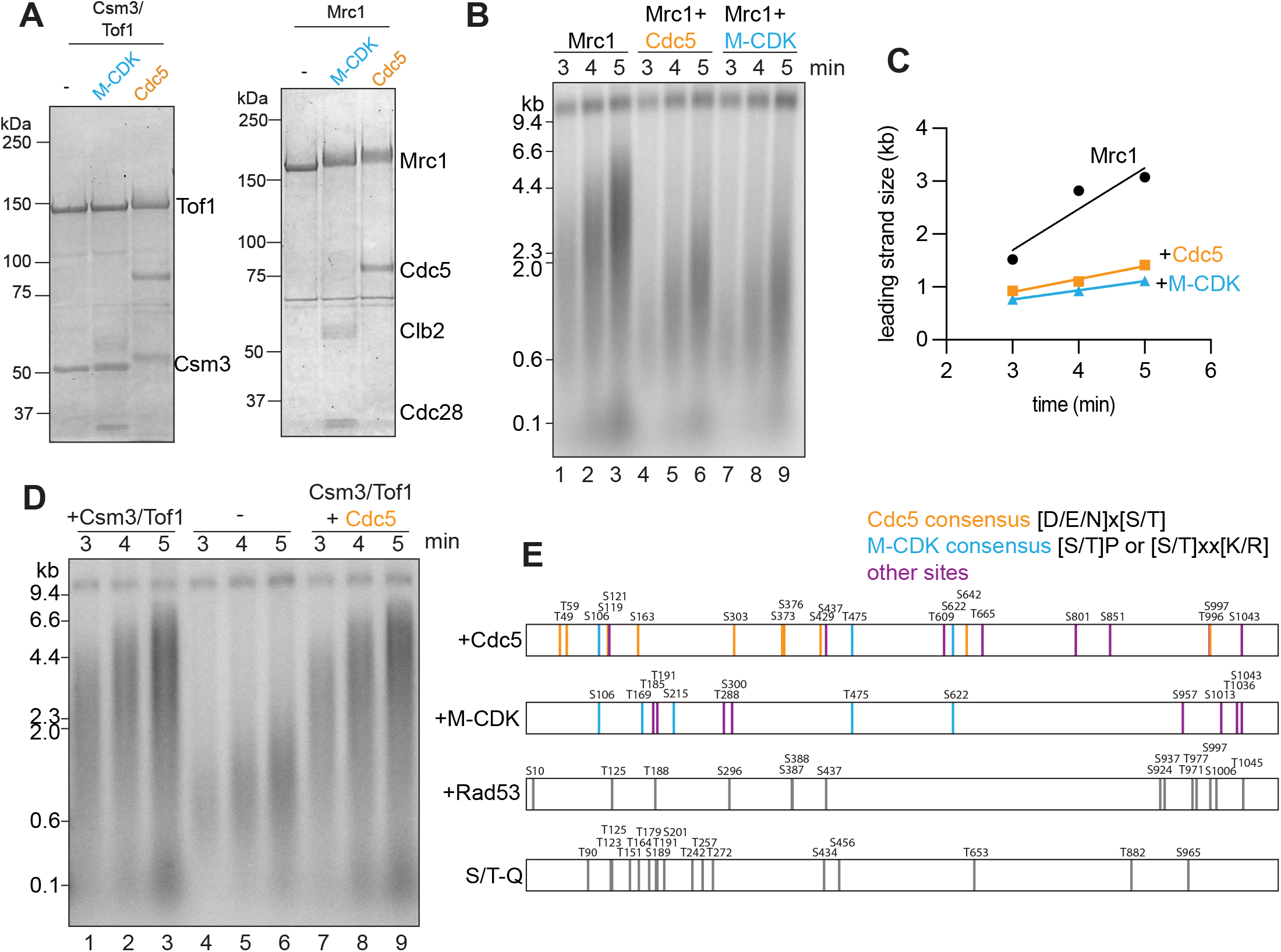
Mitotic kinases phosphorylate Mrc1 to slow DNA replication. A) Mitotic kinases were incubated with Mrc1, Csm3, and Tof1 in the presence of ATP, then run on SDS-PAGE and coomassie stained. B) Mrc1 was pre-incubated with either Cdc5 or M-CDK and then added to an otherwise normal replication reaction. C) quantification of leading strand sizes from B). D) Csm3/Tof1 was pre-incubated with Cdc5 and then added to an otherwise normal replication reaction. E) Illustration of mass spectrometry detection of Mrc1 phospho-sites. Data of Rad53-dependent sites is adapted from McClure and Diffley, 2021^28^. S/T-Q sites are potential Mec1 sites as described in Osborn and Elledge, 2003^7^.

To identify which residues in Mrc1 are phosphorylated, we performed mass spectrometry on purified Mrc1 incubated with either Cdc5 or M-CDK. With a coverage of 78-80% of Mrc1, we identified sites that were at least 40% phosphorylated in the presence of a kinase (Fig 2E). There were 12 Cdc5-dependent sites that followed the consensus sequence of [D/E/N]x[S/T], and 5 M-CDK-dependent sites that followed either the canonical consensus sequence [S/T]P or the non-canonical consensus sequence [S/T]xx[K/R]^25–27^. There were 9 additional sites for Cdc5-phosphorylated Mrc1 and 8 additional sites for M-CDK-phosphorylated Mrc1. Unlike the Mrc1 phosphorylation sites that are Rad53-dependent, which are clustered in C-terminus^28^, or the potential Mec1-dependent sites^7^, which are clustered in the N-terminus, both Cdc5 and M-CDK phosphorylation sites in Mrc1 are distributed throughout the Mrc1 protein. Consistently, Cdc5 and M-CDK caused a slower migration of the Mrc1 mutant lacking the C-terminus (Δ876-1096) (Fig S1A), while Rad53 did not^28^.

Previous proteome-wide mass spectrometry datasets identified some of the same phospho-sites enriched in mitotic samples (S609, S622, and S997)^17,29^. Other phosphorylated sites identified from mitotic cells were also present in our purified samples (S367, S605, S607, and S807)^15–17,29^, but they failed to meet our selection criteria because they were present in the non-kinase treated control peptides >40%. Since our purified Mrc1 supported fast replication elongation in the absence of mitotic kinases, these sites are likely not responsible for the replication slowing upon incubation with mitotic kinases. We next wanted to determine whether Mrc1 showed a similar change in gel-migration in mitotic cells. Yeast cells were synchronized in G1 with α-factor, then released into fresh media containing nocodazole to induce a prometaphase arrest where Cdc5 and M-CDK activity is high^30,31^. Mrc1 did not exhibit a slower gel-migration (Fig S1B) consistent with previous data^32^. Altogether, we conclude that Mrc1 is phosphorylated by Cdc5 and M-CDK in biochemical reconstitution and in mitotic cells, though further studies are needed to determine which phosphorylated residues are responsible for slowing replication.

Mrc1 interacts with Tof1 and Csm3 to form a trimeric complex at the replication fork^33,34^. Previous cross-linking mass spectrometry data mapped S300 and T322 of Mrc1 to the interaction with Tof1^35^, and since some of the phosphorylation sites we identified fall in this vicinity, we hypothesized that the interaction between Mrc1 with Tof1 or Csm3 would be disrupted by phosphorylation. To this end, we co-immunoprecipitated Mrc1 that was pre-incubated with Cdc5 or M-CDK followed by addition of Tof1 and Csm3. Tof1 and Csm3 remained bound to Mrc1 in all samples indicating that phosphorylation did not disrupt the Mrc1-Tof1-Csm3 interaction (Fig S1C).

### M-CDK phosphorylates Polα and reduces lagging strand initiation

To observe other potential mitotic kinase regulation of replication forks, we modified the reconstituted replication pause and restart protocol to include the radiolabeled dCTP even after replication restart. This ongoing labeling allowed for visualization of Okazaki fragments that are synthesized post-restart. When M-CDK was incubated with replication forks paused by topoisomerase omission, lagging strands averaged 487 bp, a marked increase from the 289 bp observed in the absence of kinase (Fig 3A). Similar lagging strand lengthening was seen in the LacR pause and IPTG restart reactions when mitotic kinases were added (Fig 3B).

**Figure 3.**
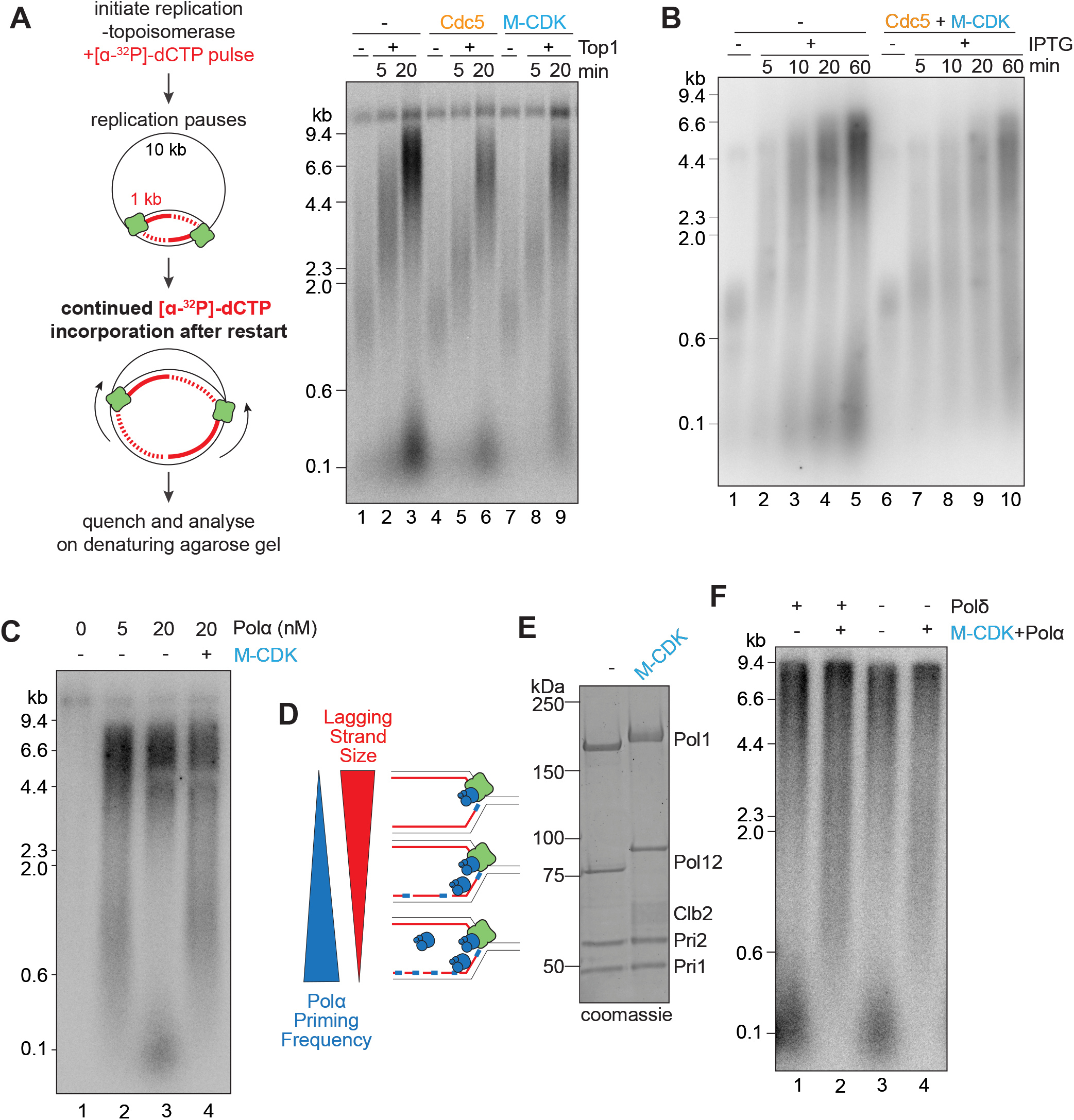
M-CDK phosphorylates Polα and reduces lagging strand initiation. A) Experimental setup like Figure 1A and 1B, altered to radiolabel DNA synthesized post replication restart. Cdc5 or M-CDK was added to replication forks paused by topoisomerase omission, then replication was restarted by the addition of TopoI. B) LacR blocked replication forks were incubated with Cdc5 and M-CDK then restarted by the addition of IPTG. C) Polα was included in normal replication assays at 0, 5 or 20 nM or pre-incubated with M-CDK. D) Model describing how lower concentrations of Polα leads to longer lagging strands due to lowered priming frequency. E) Polα was incubated with M-CDK and then run on SDS-PAGE and coomassie stained. F) Polα was incubated with M-CDK and then added to a replication reaction with or without Polδ.

Lower concentrations of Polα (the primase-polymerase responsible for lagging strand initiation) also lengthened lagging strands (Fig 3C, compare lane 2 to 3) as previously seen^20^, indicative of less frequent priming activity (Fig 3D). We therefore wondered whether M-CDK-dependent lengthening of lagging strands could be due to reduction of Polα activity. Indeed, the Pol1 and Pol12 subunits of Polα were phosphorylated by M-CDK as observed by slower gel-migration (Fig 3E) and as previously reported^36–38^. Polα pre-incubated with M-CDK also led to longer lagging strands in an otherwise normal replication reaction (Fig 3C, lane 4). We note that this data suggests that Polα activity is reduced but not completely inhibited by M-CDK phosphorylation because reactions lacking Polα produced no replication products (Fig 3C, lane 1). Following Okazaki fragment initiation by Polα, Polδ extends primers in an RFC and PCNA dependent manner^17,29^. In reactions lacking Polδ, lagging strands were still lengthened when Polα was incubated with M-CDK (Fig 3F, compare lane 4 to lane 3). Together, this suggests that M-CDK phosphorylation of Polα alone is responsible for lagging strand lengthening.

Given M-CDK regulation of lagging strand size, we next tested whether these lagging strands were still competent for nuclease processing and ligation to produce mature lagging strands. We repeated the LacR pause and restart experiment (as in Fig 3B) and included the Fen1 flap endonuclease and Cdc9 ligase. In the presence of M-CDK and Cdc5, full-length products were still visualized by 60 min indicating M-CDK-dependent lengthening of lagging strands did not affect overall lagging strand maturation (Fig S2).

### Preventing phosphorylation of Polα restores lagging strand initiation

We performed mass spectrometry on purified Polα incubated with M-CDK to identify the sites phosphorylated. We identified 12 sites on Pol1, 10 sites on Pol12, and one site on Pri2 that met our criteria (see Materials and Methods, Fig 4A). Most sites fell in unstructured regions of the proteins.

**Figure 4.**
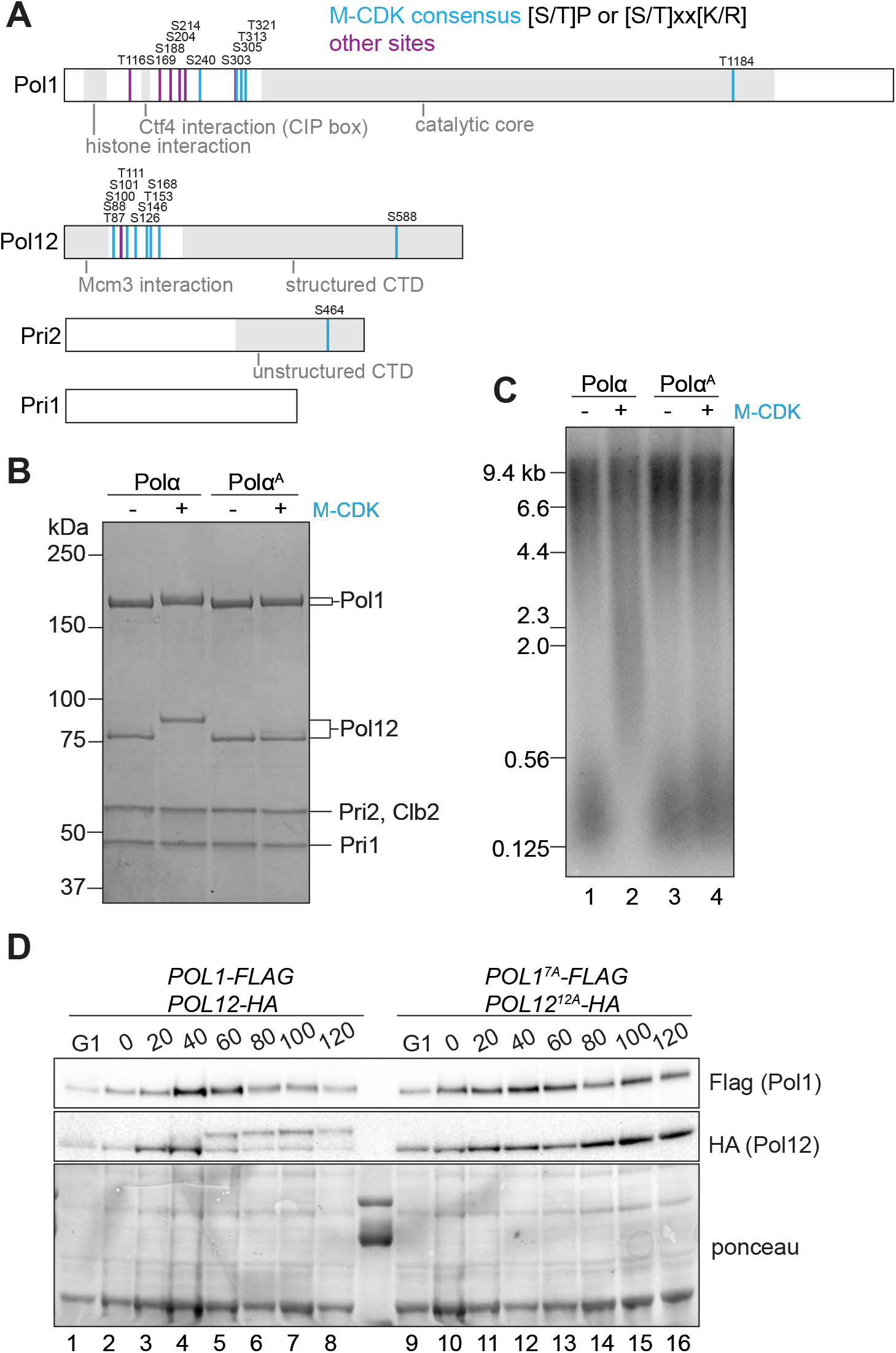
Preventing phosphorylation of Polα restores lagging strand initiation. A) Illustration of mass spectrometry detected phospho-sites on the Pol1 and Pol12 subunits of Polα. B) Polα containing the Pol1^7A^ and Pol12^12A^ mutant subunits (see Materials and Methods for list of sites) was incubated with M-CDK and run on SDS-PAGE and stained with coomassie. C) Mutant Polα from Fig 4B was added to a 20 min replication reaction. D) Cells from yAWM870 and yAWM902 were arrested in G1 with α-factor for 2.5 h, then released into media containing nocodazole. Samples were harvested at the indicated timepoints and immunoblotted with the indicated antibodies.

We purified a Polα mutant with 7 sites in Pol1 and 12 sites in Pol12 mutated to alanine, which we denote Polα^A^. The Pol1 and Pol12 subunits of Polα^A^ no longer showed shifted gel-migration in SDS-PAGE after incubation with M-CDK (Fig 4B). When Polα^A^ was included in a reconstituted replication reaction, the leading and lagging strand products were the same as reactions with wild-type Polα, but the lagging strands did not lengthen when Polα^A^ was pre-incubated with M-CDK (Fig 4C). This data suggests that the Polα^A^ is resistant to phosphorylation regulation by M-CDK. We attempted to make a mutant that would mimic phosphorylation by mutating the same residues to glutamic acids, but this mutant produced an intermediate phenotype and was still regulated in part by M-CDK, making it difficult to make meaningful conclusions (Fig S3A,B).

Pol12 phosphorylation is well documented in cells, beginning in late S phase and reaching full phosphorylation in metaphase consistent with Clb2 expression and M-CDK activity^36,37^ (Fig 4D). To determine whether Polα^A^ was resistant to phosphorylation in cells, we released cells from an α-factor G1 arrest into nocodazole to arrest them in prometaphase. Cells harboring Polα^A^ did not demonstrate slower gel-migration of Pol12 visualized by immunoblotting consistent with Polα^A^ being resistant to phosphorylation in cells (Fig 4D). The Pol1 subunit shows very little shift in gel-migration from S phase to mitosis, making it difficult to assess any changes between wild-type Pol1 and the Pol1^7A^.

Since our biochemical reconstitution data suggests that Polα phosphorylation reduces its activity at replication forks, we hypothesized that phosphorylation could reduce its affinity for replication forks and be removed from chromatin in mitotic cells. To test this, we performed fractionation by chromatin pelleting over sucrose cushions, isolating chromatin-associated proteins from soluble proteins. Consistent with previous work^36^, we found that phosphorylated Pol12 was less associated with chromatin in mitosis, but the phospho-resistant mutant Pol12^12A^ was still associated with chromatin even in mitosis (Fig S3C).

### Phosphorylation of other replication elongation proteins

To test whether M-CDK and Cdc5 might target additional replication proteins, we performed a pause and restart of replication as in Figure 1 by topoisomerase omission, but here we omitted Mrc1 and used the Polα^A^ mutant to remove the regulation we have already described. In these reactions, the addition of M-CDK and Cdc5 resulted in slower replication forks and less complete replication products (Fig 5A,B). This suggests that other replication proteins are targeted by the mitotic kinases.

**Figure 5.**
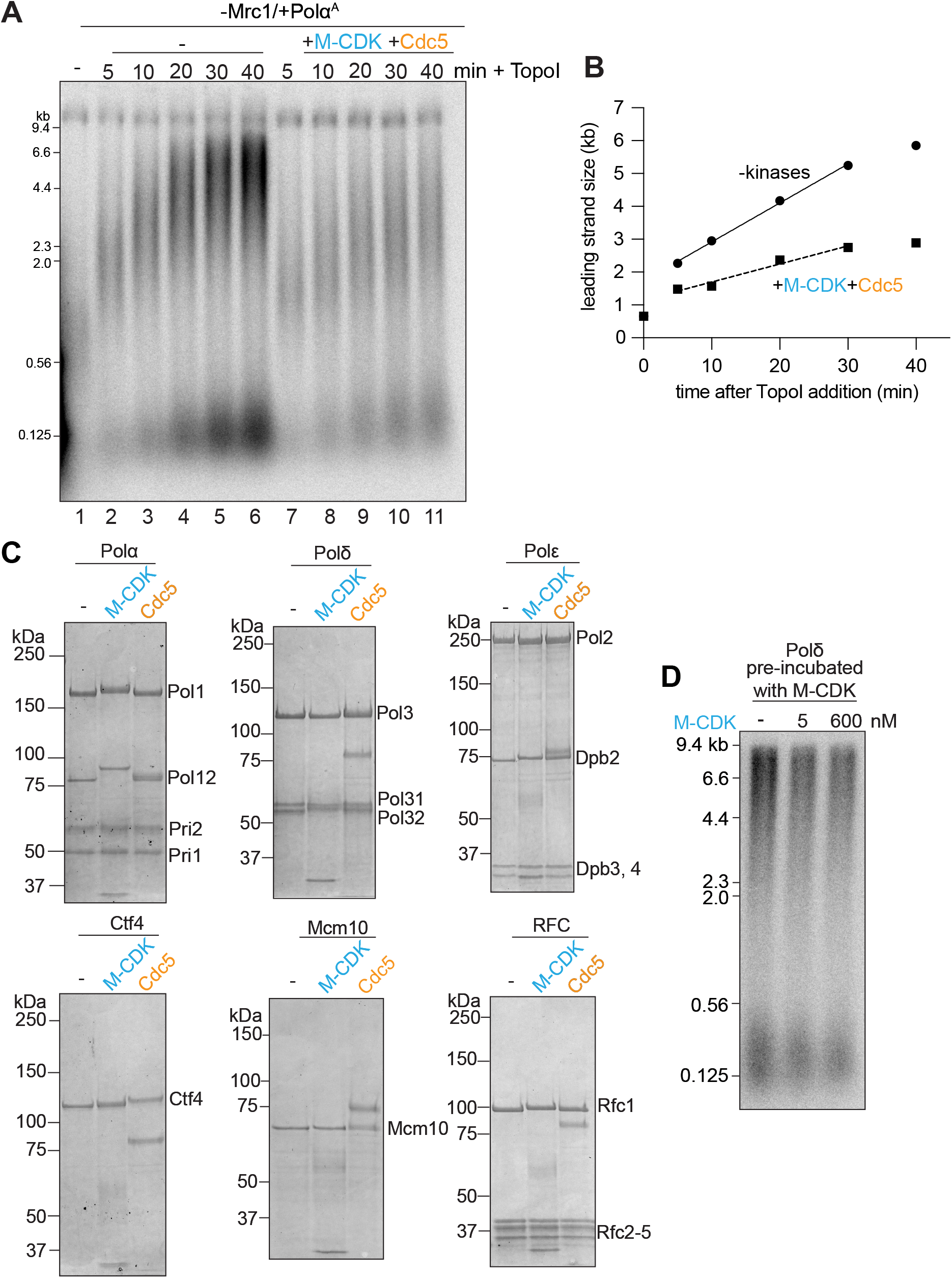
M-CDK and Cdc5 targeting of elongation replication fork proteins. A) Reconstituted replication forks were paused by topoisomerase omission in reactions lacking Mrc1 and with Polα^A^. Forks were then restarted by addition of TopoI. B) quantification of A). C) Individual replication fork elongation proteins were incubated 1:1 with either M-CDK or Cdc5 and then separated on SDS-PAGE and coomassie stained. D) Polδ was pre-incubated with 0, 5, or 600 nM M-CDK prior to addition to a 20 min replication reaction.

We then incubated replication elongation proteins individually with either M-CDK or Cdc5 and assessed potential phosphorylation by slower gel-migration. We identified subtle shifts in Pol12 in the presence of Cdc5, Pol32 in the presence of M-CDK, Dpb2 in the presence of M-CDK and Cdc5, Ctf4 in the presence of Cdc5, Mcm10 in the presence of Cdc5, and Rfc1 in the presence of M-CDK (Fig 5C). Other elongation proteins did not show any changes in gel-migration (Fig S4B). Pol32 phosphorylation by M-CDK has been previously identified^14,16^, so we tested whether this phosphorylation altered Polδ activity during replication. There were no changes to lagging strand size or amount when Polδ was pre-incubated with M-CDK suggesting phosphorylation of Pol32 does not affect Polδ activity (Fig 5D).

### Phosphorylation of Polα regulates mitotic progression

Since Polα phosphorylation is M-CDK-dependent and occurs in early mitosis, we tested whether mitotic progression might be altered in cells harboring both phospho-resistant alleles, *POL12*^*12A*^ and *POL1*^*7A*^ (*POLα*^*A*^). When we constructed the strains by tetrad dissection from heterozygous diploids, we found that colonies resulting from mutant spores were larger than wild-type colonies (Fig 6A). We then tested whether this larger colony size was a reflection of increased growth rate and found that was indeed the case (Fig 6B) with mutant cells exhibiting a doubling time of 65 min compared to wild-type cells with a doubling time of 90 min (Fig 6C).

**Figure 6.**
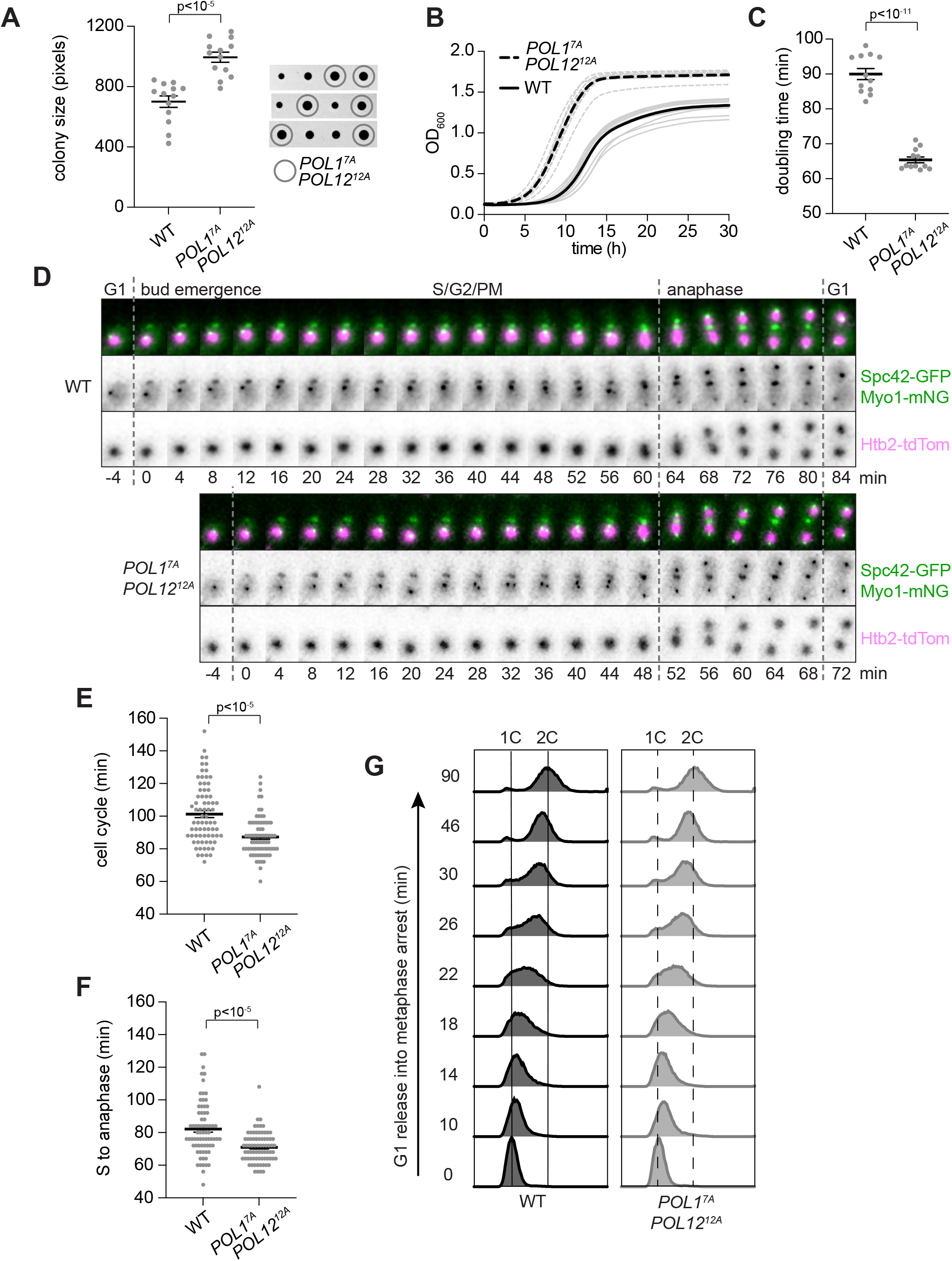
Phosphorylation of Polα regulates cell cycle progression. A) Tetrad spores from heterozygous *POL1*^*7A*^ and *POL12*^*12A*^ diploid (yAWM844) were grown for two days and then measured. B) Growth rates of 12 colonies from A) were measured in liquid culture by OD measurements every 20 min. C) Doubling time of growth rates from B). D) Live-cell imaging every 4 min was used to visualize the indicated markers. E) Cell cycle time from imaging in D) was quantified as the time between Myo1-mNG disappearance (cell division) events. F) S to anaphase timing from imaging in D) was determined by Myo1-mNG appearance to SPB (Spc42-GFP) separation and nuclear (Htb2-tdTom) entrance into the bud.

To test whether this phenotype could be seen in single cells, we performed live-cell fluorescence imaging. We tagged the histone Htb2 with tdTomato to identify the nucleus and the myosin Myo1 with mNeonGreen to identify the end of cell division (when Myo1-mNG disappears). We also tagged Spc42 with GFP to visualize the spindle, which produced distinct spots from the Myo1-mNG localization at the cell cortex. We imaged asynchronous cells at 4 min intervals and then measured overall cell cycles, defined by Myo1-mNG disappearance to the following Myo1-mNG disappearance. The *Polα*^*A*^ again exhibited faster cell cycles of 87 min compared to wild-type cells which cycled at 101 min (Fig 6E). We note that these cycles are longer than those found in the bulk liquid growth assays, which is likely due to phototoxicity^39^ and the presence of the Htb2 tag^40^.

Myo1-mNG appearance in unbudded cells marks the G1-S transition, and anaphase can be visualized by Spc42-GFP separation concurrent with nuclear (Htb2-tdTomato) extension into the bud. We found that *Polα*^*A*^ mutants spent 71 min between the onset of S phase and the onset of anaphase while wild-type cells spent 82 min (Fig 6D, F). This time encompasses S, G2, and early mitosis phases, and our imaging could not distinguish between them. To test whether S phase, defined by the time spent replicating DNA, was faster in the mutants, we performed FACS analysis of DNA content of cells first arrested in G1 with α-factor pheromone and then released into media containing nocodazole to arrest in prometaphase with 2C DNA content. *Polα*^*A*^ mutants showed similar S phase progression to wild-type cells (Fig 6G), though it should be noted that bulk DNA replication by FACS reflects both the number of DNA replication forks and replication speed^41^.

Since chromosome segregation in anaphase immediately follows metaphase where Polα phosphorylation peaks, we examined whether chromosome segregation timing was altered in the *Polα*^*A*^ mutants. In our live cell movies, late mitosis, which we define as the initiation of anaphase visualized by SPB separation and nuclear extension into the bud to cell division visualized by Myo1-mNG disappearance was 20 min for both WT and *Polα*^*A*^ mutants (Fig 6D). We also performed shorter live-cell confocal imaging for higher spatial and time resolution (2’ intervals). *Polα*^*A*^ mutants again exhibited the same late mitosis timing as wild-type cells (20 min, Fig S5A,B).

Altogether, we conclude that the *Polα*^*A*^ mutant exhibits faster cell cycles likely due to a faster early mitotic phase (prophase and metaphase), which is consistent with Polα phosphorylation occurring in early mitosis concomitant with the expression of Clb2 and rise of M-CDK activity. Therefore, this data suggests that Polα phosphorylation by M-CDK signals to the cell cycle machinery to delay mitotic progression.

## Discussion

### Mitotic remodeling of S phase replication forks

We demonstrate that the mitotic kinases M-CDK and Cdc5 regulate many proteins at DNA replication forks. We characterize two replication elongation proteins, Mrc1 and Polα, whose phosphorylation by mitotic kinases leads to their inhibition. Phosphorylation of Mrc1 results in slower replication and phosphorylation of Polα reduces lagging strand initiation.

Why would a cell entering mitosis actively inhibit replication forks instead of accelerating their completion before chromosome segregation? We speculate that this inhibition serves as a regulatory switch, remodeling replication forks to transition from canonical replication to MiDAS. MiDAS in mammalian cells requires Rad52, Mus81, and POLD3 that are dispensable for S phase replication but are required for repair processes like Break-Induced Replication (BIR)^1,42–47^. This transition would likely require removal of the S phase replisome including CMG disassembly, and phosphorylation of Mrc1 and Polα could be involved. For example, the reduced speed of replication upon Mrc1 phosphorylation could create a window of opportunity to structurally remodel the fork which may not be possible during efficient S phase replication. In addition, reduced lagging strand initiation activity by Polα could leave a persistent 3’ end of an Okazaki fragment to be processed for BIR.

In addition to our results identifying Mrc1 as a target of M-CDK and Cdc5, the DNA damage response kinases Mec1 and Rad53 also target Mrc1^7,28,32^. Biochemically reconstituted replication reactions demonstrate that phosphorylation by Cdc5, M-CDK, and Rad53 each result in a comparable reduction in replication speed, despite the kinases targeting distinct sites on Mrc1 (Fig 2). Mechanistically, Mrc1 phosphorylation may trigger dissociation of Mrc1 or modify its function at replication forks to slowdown replication. Rad53 phosphorylation of the C-terminus of Mrc1 did not result in dissociation from replication forks^28^, but since Cdc5 and M-CDK phosphorylated other regions of Mrc1, it remains unclear whether their phosphorylation would result in full dissociation. Alternatively, this slowdown may be a secondary effect of Mrc1 phosphorylation that primarily promotes another function related to replication fork protection and remodeling. For example, Mrc1 phosphorylation could promote recruitment of the ubiquitin ligase Dia2 to facilitate Mcm7 ubiquitination and subsequent CMG disassembly^48,49^.

While M-CDK phosphorylation of the Pol12 and Pol1 subunits of Polα is well-documented^36–38^, the functional consequences of phosphorylation have been unclear. We demonstrate that M-CDK mediated phosphorylation of Polα directly impairs its lagging strand initiation activity. Much like the mitotic regulation of Mrc1, this inhibition may result from dissociation of Polα from the replisome. We mapped the phospho-sites to the intrinsically disordered regions (IDRs) of both Pol1 and Pol12, which notably remain unresolved in existing structures^50,51^. Future studies are needed to determine the precise mechanism by which phosphorylation disrupts Polα function.

Beyond canonical S phase replication, Polα interacts with the CST complex to maintain telomeres^52–54^ and has been implicated in BIR^55,56^. Polα phosphorylation could regulate its activity in these other contexts. Polα has also been reported to move to the nuclear periphery in mitosis^57^, raising the intriguing possibility that such localization may be triggered by M-CDK phosphorylation and could functionally link its roles of telomere replication and localization^58^.

Our study also found that the mitotic kinases target S phase replication fork components beyond Mrc1 and Polα. Notably, the Pol32 subunit of Polδ, whose mammalian homolog POLD3 is implicated in MiDAS and BIR^43,55^, is phosphorylated by both M-CDK and Cdc5. Since our reconstituted biochemical system only includes core S phase replication machinery, it lacks the broader proteomic context (specifically MiDAS-associated factors and additional mitotic kinases) required to fully resolve the functional impact of these phosphorylation events. Nevertheless, our findings suggest that mitotic kinases directly modify S phase replisomes, providing a robust molecular foundation for investigating mitotic replication fork regulation.

### Potential negative feedback mechanism between Polα and cell cycle progression

We find that mutating Polα M-CDK phosphorylation sites *in vivo* accelerates cell cycle progression by shortening early mitosis (Fig 6). Although disruptive mutations in essential pathways typically reduce fitness and slow growth, the shortened cell cycle and faster growth in *POLα*^*A*^ mutants may reflect the bypass of a regulatory brake or negative feedback mechanism that ordinarily slows mitotic progression. While our reconstituted biochemical system defined the immediate molecular consequences of Polα phosphorylation, this cellular phenotype suggests that Polα phosphorylation imposes a checkpoint-like delay. We speculate that the presence of phosphorylated Polα at unfinished replication forks serves as a signal to delay mitotic progression until these replication forks are processed or resolved.

This finding is particularly compelling given the absence of a known checkpoint to monitor the completion of DNA replication at the end of S phase. Traditionally, crosstalk between replication forks and cell cycle progression has focused on the S phase replication checkpoint which senses double-stranded DNA breaks or excessive single-stranded DNA^41,59–61^. An additional pathway utilizing the release of firing factors as a proxy for replication completion has been suggested^62^, but it is unlikely to detect a small number of unfinished replication forks. Here, we uncover a new pathway through which a core replisome component may signal to the cell cycle machinery. Key questions remain regarding the precise context of Polα phosphorylation and the specific regulatory information it conveys to the cell cycle, but the conservation of Polα phosphorylation in human cells^63^ suggests that these functional consequences could represent a fundamental feature of the eukaryotic cell cycle.

## Materials and Methods

### Yeast Strains and plasmids

Yeast strains were constructed with standard techniques using the W303 background^64^ and are listed in Table S1. Plasmids constructed are listed in Table S2. Construction details are listed in the Supplemental Materials and Methods.

### Protein Purification

Protein expression and purification methods are summarized in Table S3, and details are in the Supplemental Materials and Methods.

### Replication reactions

The LacR pause was completed using a modified version of the previously published protocol^65^ where 5000 nM LacR and 58 nM of plasmid pAWM197 were incubated in a 30°C thermomixer at 1250 rpm for 15 min to create the replication fork block and addition of 40 mM IPTG to release LacR from the DNA. The replication reactions were performed as previously published^19,28^. The buffer used during the replication reaction contained 25 mM HEPES-KOH pH 7.5, 20 mM magnesium acetate, 0.02% NP-40-S, 2 mM DTT, and either 100 mM or 250 mM potassium glutamate.

### Quantification of leading strand lengths

Square-root encoded images (.gel) were first transformed to linear data using the Linearize GelData plugin for ImageJ. The same rectangular ROI was selected around each lane of the gel and signal plotted using Plot Profile in Fiji. The raw values for the HindIII ladder were fitted to an exponential equation to extrapolate gel migration to kb. The maximum signal value was selected as the point of measurement for leading strand length. Product length was plotted and where applicable the rate of synthesis was calculated from the linear region of the curve.

### Kinase Assays

Kinase assays were performed in buffer containing 25 mM HEPES-KOH pH 7.5, 10 mM magnesium acetate, 100 mM potassium glutamate, 0.02% NP-40-S, 2 mM DTT, and 1 mM ATP. The target protein (200 nM) was incubated with 200 nM M-CDK and/or Cdc5. Equal volumes of M-CDK and Cdc5 storage buffer were added to samples without kinases to account for the different buffer components. The samples were incubated in a 30°C thermomixer at 1250 rpm for 15 min and visualized on either 3-8% Tris-Acetate PAGE or 4-20% TGX PAGE (Bio-Rad) and coomassie stained.

### Mass spectrometry

Samples from kinase assays were analyzed for mass spectrometry, and details of LC-MS/MS analysis are in Supplemental Materials and Methods. Phosphorylated peptides detected were converted to per amino acid basis using a custom R script and are listed in Dataset 1. Sites determined to be phosphorylated must have been present in at least 4 peptides, >40% of peptides detected phosphorylated in the sample with kinase, and <40% of peptides detected phosphorylated in the sample without kinase.

### Immunoblotting

Samples were harvested by centrifugation, then washed twice with cold PBS. Cells were then resuspended in buffer containing 2 M sorbitol, 50 mM Tris pH 8, and 2 mM MgCl^2^. Then, acid-washed glass beads and an equal volume of 20% trichloroacetic acid were added, followed by 10 min of vortexing at 4ºC. The supernatant was collected, and the beads were washed 2x with ice cold 5% TCA and added to the supernatant. Protein was then pelleted at 15,000 xg for 10 min at 4ºC, then washed twice in 1 M Tris pH 8, and resuspended in Laemmli buffer. Extracts were separated on a 3-8% SDS-PAGE and transferred to a nitrocellulose membrane, and immunoblotted with anti-HA (mouse clone 12CA5, 1:1000), or anti-FLAG (mouse clone M2, 1:1000). Signal was detected following incubation with ECL solution (Thermo Fisher) using the ChemiDoc MP imaging system (BioRad).

### Cell Growth Rate and Doubling time analysis

Growth kinetics of quadruplicate microcultures in 96-well plates were monitored by optical density (OD^600^) measurements every 20 min in a Cytation3 (BioTek) plate reader heated to 30ºC. The data were fit with a logarithmic curve f(x) = x^1^+L/(1 + e^-k(x-x0)^) and solved for coefficients L, x0 and k. Doubling time was calculated using the standard formula ln(2)/k.

### Flow Cytometry Analysis (FACS)

420 µL of culture was added to 980µL pre-chilled 100% ethanol for cell fixation. RNA was digested with 0.1mg/mL RNAse at 37ºC in 50 mM Tris-HCl pH 7.5 buffer. DNA was stained by resuspending cells in 200 mM Tris-HCl pH 7.5, 210 mM NaCl, 78 mM MgCl^2^, and 0.025 mg/mL propidium iodide. Cells were sonicated and analyzed using an Agilent NovoCyte Penteon. For each sample, 20,000 events were counted and data was analyzed using FlowJo software.

### Widefield live-cell imaging

Mid-log phase cells grown in CSM were adhered to 35-mm glass-bottom culture dishes (MatTek Corporation) coated with 0.2 mg/mL Concanavalin A and overlaid with fresh CSM prior to imaging. Time-lapse images were acquired in an environmental chamber maintained at 30°C using a DeltaVision Ultra microscope equipped with a 60×/1.42 NA objective, immersion oil, an EDGE sCMOS camera, and a Polychroic B-G-R-FR filter set. Images were acquired every 4 minutes for 3.5 hours across 31–35 z-sections collected at 0.2 µm intervals. Exposure settings were 0.01 s at 2% illumination power for the green channel, 0.015 s at 2% illumination power for the red channel, and 0.05 s at 10% illumination power for differential interference contrast (DIC) reference images. Acquired images were deconvolved using enhanced ratio settings, and z-stacks were maximum-intensity projected in SoftWorx (GE Healthcare Life Sciences, version 7.2.2). Images were manually analyzed. Only parental cells were scored. Entry into S phase was defined by the appearance of the Myo1-mNG ring. Mitotic entry was defined as spindle pole separation. Anaphase onset was defined as the point at which spindle poles and segregating chromatin passed through the Myo1-mNG ring. Mitotic exit was defined by contraction and disappearance of the Myo1-mNG ring. All imaging experiments were performed in three independent biological replicates. For display in Fig 6, max projections of the z-series were generated and then denoised in FIJI. Intensity and color adjustments were identical for all strains and processed in FIJI.

### Declaration of generative AI and AI-assisted technologies in the writing process

During the preparation of this work, ChatGPT and Gemini were used for improving language and clarity of the writing. The authors reviewed and edited the content and take full responsibility for the content of the manuscript.

## Acknowledgements

We thank Michael McCarron and Jenna Kotz for media prep and assistance with strain construction. We thank Jeff Moore, Michael McMurray, and Julie Cooper for strains, plasmids, and use of equipment. We thank Rahul Thadani, Rishi Nageshan, the Heasley Lab, and the DDR supergroup for stimulating discussions. We thank David McClure for custom R scripts. This work was funded by the NIH grant R35GM156773 (A.W.M.) and T32GM136444 (S.R.G.). This study was supported in part by the NIH P30CA06934 funded Mass Spectrometry Proteomics Shared Resource and Flow Cytometry Shared Resource.

## Author Contributions

S.S.: Conceptualization, Formal analysis, Investigation, Methodology, Project administration, Visualization, Writing - Original Draft, Writing - Review & Editing; N.P.: Formal analysis, Investigation, Methodology, Visualization, Writing - Review & Editing; A.M.: Investigation, Methodology, Writing - Review & Editing; S.N.S.: Formal analysis, Investigation, Methodology, Writing - Review & Editing; S.R.G.: Formal analysis, Investigation, Methodology; S.B.: Investigation; A.W.M.: Conceptualization, Formal analysis, Funding acquisition, Investigation, Methodology, Project administration, Supervision, Visualization, Writing - Original Draft, Writing - Review & Editing.

