## Supplemental Material for "Mitotic kinase regulation of DNA replication forks"

**Supplemental Materials and Methods**

_Yeast Strains and plasmids_

Yeast strains were constructed with standard techniques using the W303 background[1] and are listed in Table S1. Plasmids constructed are listed in Table S2.

pAWM201 containing *POL1*$^{7A}$ was made by Gibson assembly with a synthesized fragment from Twist Bioscience and verified by full-plasmid sequencing. *POL1*$^{7A}$ contains the following amino acids mutated to alanine: S240, T313, S214, S169, S305, S188, and T321. pAWM201 was transformed into a diploid and verified for proper integration with sequencing.

pAWM216 containing *POL12*$^{12A}$ was made by Gibson assembly with a synthesized fragment from Twist Bioscience and verified by full-plasmid sequencing. *POL12*$^{12A}$ contains the following amino acids mutated to alanine: S101, T111, T153, T177, T185, T190, T87, S88, S126, S168, S146, and S100. pAWM216 was transformed into a diploid and verified for proper integration with sequencing.

pAWM147 was made by Gibson assembly with codon-optimized *CLB2* synthesized fragments from Twist Bioscience. Products were verified by full-plasmid sequencing.

pAWM218 and pAWM222 were made by Gibson assembly with codon-optimized *POL1* and *POL12* synthesized fragments from Twist Bioscience. Products were verified by full-plasmid sequencing.

*POL1-3xFLAG* was made by transforming PCR products from template *pYM-3xFlag-nat-NT2* into a *POL*$^{7A}$*/POL1* diploid. Integration to *POL1* or *POL1*$^{7A}$ was determined by PCR.

*POL12-3xHA* was made by transforming PCR products from template *pFA6a-3xHA-kanMX6*[2] into a *POL12*$^{12A}$*/POL12* diploid. Integration to *POL12* or *POL12*$^{12A}$ was determined by PCR.

PCR based tagging methods using pFA6 plasmids were utilized to create *MYO1-mNeonGreen:kan*$^{R}$ and *SPC42-GFP:kan*$^{R}$ (Table S1). *HTB2-tdTomato:HIS3* was created by PCR from a strain containing *HTB2-tdTomato:HIS3*[4].

Protein purification strains were essentially the same as previously used[3]. The strains were modified with the β-estradiol-sensitive hybrid transcription factor *GAL4*$^{DBD}$*-hER-VP16*$^{AD}$. Updated genotypes for these strains are listed in Table S1. pAWM205 was made by modifying pPP1557[5] with the pRS40N[6] vector using ApaI and SacII. The product was verified by full-plasmid sequencing. The plasmid was cut using AhdI to target genome integration (into a strain that already contained *amp*$^{R}$ cassette).

**Table S1.**

| ***yeast strain*** | ***genotype*** | ***source*** |
|---|---|---|
| yAWM336 | *MATa*<br>*MRC1-3xFLAG:nat*$^{R}$ | McClure and Diffley, 2021 |
| yAWM844 | *MATa/α*<br>*ura3:POL1*$^{7A}$*:URA3/POL1*<br>*leu2:POL12*$^{12A}$*:LEU2/POL12* | this study |
| yAWM870 | *MATa*<br>*POL1-3xFLAG:nat*$^{R}$<br>*POL12-3xHA:kan*$^{R}$ | this study |
| yAWM882 | *MATa*<br>*his3:HTB2-tdTomato:HIS3*<br>*MYO1-mNeonGreen:kan*$^{R}$ | this study |
| yAWM880 | *MATa*<br>*his3:HTB2-tdTomato:HIS3*<br>*MYO1-mNeonGreen:kan*$^{R}$<br>*ura3:POL1*$^{7A}$*:URA3*<br>*leu2:POL12*$^{12A}$*:LEU2* | this study |
| yAWM902 | *MATa*<br>*ura3:POL1*$^{7A}$*-3xFLAG:URA3:nat*$^{R}$<br>*leu2:POL12*$^{12A}$*-3xHA:LEU2:kan*$^{R}$ | this study |
| yAWM900 | *MATα*<br>*SPC42-GFP:kan*$^{R}$<br>*his3:HTB2-tdTomato:HIS3*<br>*MYO1-mNeonGreen:kan*$^{R}$ | this study |
| yAWM901 | *MATα*<br>*ura3:POL1*$^{7A}$*:URA3*<br>*leu2:POL12*$^{12A}$*:LEU2*<br>*his3:HTB2-tdTomato:HIS3*<br>*MYO1-NeonGreen:kan*$^{R}$<br>*Spc42-GFP:kan*$^{R}$ | this study |

| | | |
|---|---|---|
| yAWM847 | *MATa*<br>*bar1::hyg$^R$*<br>*pep4::kan$^R$*<br>*trp1:Gal1-10 ORC5, ORC6:TRP1*<br>*his3:Gal1-10 ORC3, ORC4:HIS3*<br>*ura3:Gal1-10 CBP-ORC1, ORC2:URA3*<br>*GEV::nat$^R$* | this study |
| yAWM851 | *MATa*<br>*bar1::hyg$^R$*<br>*pep4::kan$^R$*<br>*his3:pRS303/CDT1, GAL4:HIS3*<br>*trp1:pRS304/MCM4, MCM5:TRP1*<br>*leu2:pRS305/MCM6, MCM7:LEU2*<br>*ura3:pRS206/MCM2, FLAG-MCM3:URA3*<br>*GEV::nat$^R$* | this study |
| yAWM848 | *MATa*<br>*pep4::kan$^R$*<br>*trp1:pRS304CDC7, CBP-DBF4:TRP1*<br>*GEV::nat$^R$* | this study |
| yAWM839 | *MATa*<br>*bar1::hyg$^R$*<br>*pep4::kan$^R$*<br>*his3:pRS303/Dpb11-3Xflag (Nat-NT2), GAL4:HIS3*<br>*trp1:GEV:TRP1* | this study |
| yAWM852 | *MATa*<br>*bar1::hyg$^R$*<br>*pep4::kan$^R$*<br>*ura3:pRS306/Dpb2, Dpb3:URA3*<br>*trp1:pRS304/Pol2, Dpb4-Tev-CBP:TRP1*<br>*GEV::nat$^R$* | this study |

| | | |
|---|---|---|
| yAWM840 | *MATa*<br>*bar1::hyg$^R$*<br>*pep4::kan$^R$*<br>*ura3:Gal1-10 CKS1+CDC28:URA3*<br>*his3:Gal1-10 CBP-Tev-Clb5Δ1-100+Gal4:HIS3*<br>*trp1:GEV:TRP1* | this study |
| yAWM696 | *MATa*<br>*bar1::hyg$^R$*<br>*pep4::kan$^R$*<br>*his3:pRS303-Sld3sup-TCPsup:HIS3*<br>*leu2:pRS305-Sld7sup:LEU2*<br>*trp1:GEV:TRP1* | this study |
| yAWM647 | *MATa*<br>*bar1::hyg$^R$*<br>*pep4::kan$^R$*<br>*his3:Gal-Mrc1-2xflag:HIS3*<br>*trp1:GEV:TRP1* | this study |
| yAWM841 | *MATa*<br>*bar1::hyg$^R$*<br>*pep4::kan$^R$*<br>*his3:pRS303/CBP-Tev-RFA1, GAL4:HIS3*<br>*ura3:pRS306/RFA2, RFA3:URA3*<br>*trp1:GEV:TRP1* | this study |
| yAWM842 | *MATa*<br>*bar1::hyg$^R$*<br>*pep4::kan$^R$*<br>*his3:pRS303/CBP-Tev-Ctf4, GAL4:HIS3*<br>*trp1:GEV:TRP1* | this study |
| yAWM853 | *MATa*<br>*bar1::hyg$^R$*<br>*pep4::kan$^R$*<br>*his3:pRS303-CBP-TOP1+Gal4:HIS3*<br>*trp1:GEV:TRP1* | this study |

| | | |
|---|---|---|
| yAWM838 | *MATa*<br>*bar1::hyg*$^R$<br>*pep4::kan*$^R$<br>*ura3:pRS306-CBP-Csm3 + Tof1:URA3*<br>*trp1:GEV:TRP1* | this study |
| yAWM837 | *MATa*<br>*pep4::kan*$^R$<br>*ura3:pRS306-POL31+POL3:URA3*<br>*his3:pRS303-Pol32-CBP+Gal4:HIS3*<br>*trp1:GEV:TRP1* | this study |
| yAWM850 | *MATa*<br>*bar1::hyg*$^R$<br>*pep4::kan*$^R$<br>*trp1:pRS304/Pol1/Pol12:TRP1*<br>*ura3:pRS306/CBP-Tev-Pri1/Pri2:URA3*<br>*GEV::nat*$^R$ | this study |
| yAWM600 | *MATa*<br>*bar1::hyg*$^R$<br>*pep4::kan*$^R$<br>*his3:pRS303-Gal4-Gal1/10-CDC5:HIS3*<br>*trp1:GEV:TRP1*<br>*cdc28-as1* | this study |
| yAWM871 | *MATa*<br>*bar1::hyg*$^R$<br>*pep4::kan*$^R$<br>*ura3:pRS306-CKS1,Gal1-10,CDC28:URA3*<br>*his3:pRS303-Gal4-Gal1/10-Clb2*$^{\Delta 1-34}$*-CBP-TEV:HIS3*<br>*trp1:GEV:TRP1* | this study |
| yAWM843 | *MATa*<br>*pep4::kan*$^R$<br>*Pol12-3HA:kan*$^R$<br>*ura3:pRS306-Gal-Pri1,Pri2:URA3*<br>*trp1:pRS304-Pol1-7A-Gal1,10-Pol12-12A:TRP1*<br>*GEV:nat*$^R$ | this study |

**Table S2. Plasmids**

| *plasmid* | *genotype* | *source* |
|---|---|---|
| pBP83 | pYM-3Flag-nat-NT2 | Frigola, et al 2013 |
| pPP1557 | pRS304-pADH1-[GAL4DBD-hER-VP16] “GEV:TRP1” | Louvion, et al, 1993 and Takahashi and Pryciak, 2008 |
| pAWM117 | pFA6a-3HA-kanMX6 | Bahler, et al, 1998 |
| pAWM138 | pRS40N-natMX4 | Chee and Haase, 2012 |
| pAWM147 | pRS303-Gal4-Gal1/10-Clb2(Δ1-34)-CBP-TEV | this study |
| pAWM201 | pRSII306-Pol1-7A (Δ1488-4407bp) | this study |
| pAWM205 | pRS40N-$P_{ADH1}$-$GAL4^{DBD}$-hER-$VP16^{AD}$:$nat^{R}$ “GEV:$nat^{R}$” | this study |
| pAWM216 | pRS30-$POL12^{12A}$(Δ1-136bp) | this study |
| pAWM197 | p177bp-LacOx16-(artificial 70bp ARS)-LacOx19 | this study |
| pAWM174 | pFA6a-mNeonGreen:KANMx | gift from Jeff Moore, modified from Sheff and Thorn, 2004 |
| pJY22 | 7.7 kb Ars1 in Bluescript KS+ | Yeeles, et al, 2017 |
| pAWM218 | pRS304-Pol1-7A-Gal1/10-Pol12-12A | this study |
| pAWM222 | pRS304-Pol1-7E-Gal1/10-Pol12-12E | this study |

*Protein Purification*

Protein expression and purification methods are summarized in Table S3. Proteins expressed in yeast were either induced by the addition of 2% galactose or 150 nM of β-estradiol to a log phase culture or following arrest with nocodazole or α-factor. The proteins were expressed for 2-4 h then harvested at 12,000 x g for 10 min. The pellets were resuspended in lysis buffer, frozen in liquid nitrogen, and lysed in a freezer mill.

Proteins expressed in bacteria were induced by the addition of 1 mM IPTG (or 0.1 g/mL arabinose for the LacR) at log phase. The cultures were expressed then harvested at 12,000 x g for 10 min. The pellets were stored at -80°C until purified. Cells were lysed by sonication.

For the expression and purification of M-CDK, yeast cells were arrested with nocodazole at log phase and induced with 150 nM β-estradiol. After 2 h, cells were harvested at 12,000 x g for 10 min. Pellets were resuspended with 0.4 volumes of buffer containing 25 mM Tris-HCl pH 7.5, 400 mM NaCl, 0.02% NP-40-S 1 mM DTT, 10%

glycerol, and 3X Halt Protease Inhibitor Cocktail EDTA-free (Thermo Fisher) then frozen in liquid nitrogen followed by lysis in a freezer mill. The lysate was centrifuged in a Ti45 rotor (Beckman) at 45,000 rpm for 45 min at 4°C and the lysate was collected and rotated with calmodulin resin with 2 mM calcium chloride at 4°C for 1 h. An on-column TEV cleavage was performed at 4 C overnight. The subsequent eluate was loaded onto Capto S (Cytiva) column with buffer containing 25 mM Tris-HCl pH 7.5, 1 mM DTT, 0.02% NP-40-S and 10% glycerol and eluted with a 150 mM potassium acetate to 1.2 M potassium acetate gradient over 15 CV. Pooled fractions were combined and run over an S200 column with the storage buffer containing 40 mM HEPES-KOH pH 7.5, 300 potassium acetate, 1 mM DTT, 0.02% NP-40-S and 10% glycerol. Protein concentration was determined by Detergent Compatible Bradford Assay (Pierce).

For the expression and purification of Cdc5, yeast cells were arrested with nocodazole at log phase and induced with 150 nM β -estradiol. After 3 h, cells were harvested at 12,000 x g for 10 min. Pellets were resuspended with 0.4 volumes of buffer containing 45 mM HEPES-KOH pH 7.6, 300 mM potassium acetate, 0.02% NP-40-S 1 mM DTT, 1 mM EDTA ,10% glycerol, and 3X Halt Protease Inhibitor Cocktail EDTA-free (Thermo Fisher) then frozen in liquid nitrogen followed by lysis in a freezer mill. The lysate was centrifuged in a Ti45 rotor (Beckman) at 45,000 rpm for 45 min at 4°C and the lysate was collected and rotated with Anti-Flag resin for 2 h at 4°C. The protein was eluted with 0.5 mg/mL 3X flag peptide (GenScript). The eluate was loaded onto Capto Q (Cytiva) column with buffer containing 45 mM HEPES-KOH pH 7.6, 0.02% NP-40-S 1 mM EDTA, and 10% glycerol and eluted with a 100 mM potassium acetate to 1 M potassium acetate gradient over 10 CV. The protein was dialyzed for a total of 2 h at 4°C in 1 L of buffer containing 45 mM HEPES-KOH pH 7.6, 150 mM potassium acetate, 0.02% NP-40-S 1 mM DTT, and 40% glycerol with a change into fresh buffer after 1 h. Protein concentration was determined by Detergent Compatible Bradford Assay (Pierce).

**Table S3. Protein purification strategy**

| **Protein** | **Purification strategy (see [7–10] for more details)** |
|---|---|
| **MCM-Cdt1** | Yeast Expression, Calmodulin Pull-Down, EGTA Elution, Gel Filtration |
| **Cdc6** | Bacterial Expression, Glutathione Pull-Down, PreScission Protease Elution, HTP Column, Dialysis |
| **ORC** | Yeast Expression, Calmodulin Pull-Down, EGTA Elution, Gel Filtration |
| **DDK** | Yeast Expression, Calmodulin Pull-Down, EGTA Elution, Gel Filtration |
| **Mrc1** | Yeast Expression, Flag Pull-Down, Flag Peptide Elution, MonoQ, Dialysis |

| | |
|---|---|
| **Cdc45** | Yeast Expression, Flag Pull-Down, Flag Peptide Elution, HTP Column, Dialysis |
| **Dpb11** | Yeast Expression, Flag Pull-Down, Flag Peptide Elution, Gel Filtration |
| **Polε** | Yeast Expression, Calmodulin Pull-Down, EGTA Elution, Heparin Column, Gel Filtration |
| **GINS** | Bacterial Expression, Ni-NTA Pull-Down, Imidazole Elution, MonoQ, Gel Filtration |
| **CDK** | Yeast Expression, Calmodulin Pull-Down, TEV Elution, Ni-NTA Column, Gel Filtration |
| **RPA** | Yeast Expression, Calmodulin Pull-Down, EGTA Elution, Heparin Column, Gel Filtration |
| **Ctf4** | Yeast Expression, Calmodulin Pull-Down, EGTA Elution, MonoQ, Gel Filtration |
| **TopoI** | Yeast Expression, Calmodulin Pull-Down, EGTA Elution, Gel Filtration |
| **Csm3/Tof1** | Yeast Expression, Calmodulin Pull-Down, TEV Elution, Gel Filtration |
| **Polα and all Polα mutants** | Yeast Expression, Calmodulin Pull-Down, EGTA Elution, MonoQ, Gel Filtration |
| **Sld3/7** | Yeast Expression, IgG Sepharose 6 Pull-Down, TEV Elution, Ni-NTA Column, Gel Filtration |
| **Mcm10** | Bacterial Expression, Ni-NTA Pull-Down, Imidazole Elution, Gel Filtration |
| **Sld2** | Yeast Expression, Ammonium Sulfate Precipitation, Flag Pull-Down, Flag Peptide Elution, SP Column, Dialysis |
| **RFC** | Yeast Expression, Calmodulin Pull-Down, EGTA Elution, MonoS, Gel Filtration |
| **PCNA** | Bacterial Expression, Ammonium Sulfate Precipitation, SP Column, Heparin Column, DEAE Column, MonoQ, Gel Filtration |
| **Polδ** | Yeast Expression, Calmodulin Pull-Down, EGTA Elution, Heparin Column, Gel Filtration |
| **Cdc5** | Yeast Expression, Flag Pull-Down, Flag Peptide Elution, Capto Q, Dialysis |
| **M-CDK** | Yeast Expression, Calmodulin Pull-down, TEV Elution, SP Column, Gel Filtration |
| **LacR** | Bacterial Expression, Ni-NTA Pull-Down, Gel Filtration |
| **Cdc9** | Yeast Expression, Flag Pull-Down, Flag Peptide Elution, Gel Filtration |
| **Fen1** | Yeast Expression, Flag Pull-Down, Flag Peptide Elution, Gel Filtration |

*Chromatin pellet fractionation*

The chromatin pellet protocol was adapted from previously described protocols[11]. Briefly, log phase yeast strains were synchronized for 2 h using 10 ug/mL alpha factor to arrest cells in G1, then released into YPD + 5 ug/mL nocodazole. The S phase sample was taken at 35 min post-release. The prometaphase sample was taken at 120 min post-release. 12.5 OD units of cells were resuspended in cold 3 mL of CP1 buffer (100 mM PIPES-KOH pH 9.4, 10 mM DTT, 0.1% sodium azide, 1X Halt Protease Inhibitor Cocktail EDTA-free (Thermo Fisher), 1X Phosphatase Inhibitor cocktail (5 mM NaF, 1 mM sodium pyrophosphate and 1 mM sodium beta-glycerophosphate pentahydrate)). Cells were resuspended in 2 mL CP2 buffer (50 mM potassium phosphate pH 7.4, 600 mM sorbitol, 10 mM DTT, 1X Halt Protease Inhibitor Cocktail EDTA-free, 2 mM PMSF, 1 µM Pepstatin A, 10 µM leupeptin, 1X Phosphatase Inhibitor cocktail). Cells in each sample were spheroplasted by addition of 25 uL 20 kU/mL lyticase and incubated at 37°C for 15 min. Spheroplastation was monitored using a spectrophotometer and halted when the OD of a 1:20 dilution of the cell suspension in water was less than 10% of the value before addition of enzyme. Cells were washed with 1 mL CP3 (50 mM HEPES-KOH pH 7.5, 400 mM sorbitol, 100 mM KCl, 2.5 mM MgCl2, 1X Halt Protease Inhibitor Cocktail EDTA-free, 1X Phosphatase Inhibitor cocktail) then resuspended in 130 uL EB buffer (50 mM HEPES-KOH pH 7.5, 100 mM KCl, 2.5 mM MgCl2, 1 mM DTT, 1X Halt Protease Inhibitor Cocktail EDTA-free, 2 mM PMSF, 1 µM Pepstatin A, 10 µM leupeptin,1X Phosphatase Inhibitor cocktail).

Cells were lysed by addition of Triton X-100 to a final concentration of 0.5%, and incubated on ice for three min with occasional vortexing. Samples were centrifuged (4000 rpm, 1 min, 4ºC), and 25 uL of the supernatant whole cell extract (WCE) was removed for immunoblotting. Samples were boiled immediately after addition of 1X sample buffer. A sucrose cushion was prepared using EBX-S buffer (1X EB, 0.25% Triton X-100, and 30% sucrose). The remaining 100 uL of WCE was gently laid upon 100 uL of the EBX-S cushion, then centrifuged (12000 rpm, 20 min, 4ºC). From the top layer, 40uL is taken for immunoblotting (SUP). The remaining supernatant and sucrose layer were aspirated, and the pellet was with 300 uL EBX buffer (1X EB, 0.25% Triton X-100), then resuspended in 100 uL EBX buffer with addition of 28 units of benzonase. Sample was incubated on ice for 15 min with occasional vortexing, then 40 uL of this chromatin pellet fraction was taken for immunoblotting (CP).

*Mass spectrometry*

For LC-MS/MS analysis, 10 µg of phosphorylated protein was proteolytically digested by Trypsin (Promega) into peptides according to the suspension trap (S-Trap) digestion protocol[12]. The resulting peptides were dried down by speedvac, desalted using Pierce C18 Spin Tips (ThermoFisher) and resuspended in 0.1% formic acid (FA) for analysis. Peptide samples were analyzed using a Bruker tims-TOF SCP mass spectrometer coupled to a nanoElute nano-UHPLC (Bruker Daltronics). Peptides were separated on a PepSep 25 cm x 150 mm, C18 1.5 µm (Bruker Daltronics) heated to 50°C in a column oven. 0.1% (v/v) FA (mobile phase A) and acetonitrile (mobile phase B) were used for the separation. The separation method consisted of a 60 min stepped gradient consisting of the sequential steps: (a) 4 to 28% B over 40 min, (b) 28 to 32% B over 9.5 min, (c) 32 to 95% B over 10 min, and (d) hold at 95% B for 8 min. The flow rate for separation was 0.75 µL/min. Mass spectrometer performance was monitored by periodic injections of a K562 tryptic digest (Pierce) where the number of peptides, total ion chromatogram shape and mass calibration were considered.

The mass spectrometer was operated in PASEF mode with an accumulation time of 75 ms and 5 PASEF MS/MS scans per topN acquisition cycle. MS and MS/MS spectra were recorded for a m/z range of 100 to 1700 and ion mobility was scanned from 0.6 to 1.4 Vs/cm2. Precursors for data-dependent acquisition were isolated within ± 1 Th and fragmented with an ion mobility-dependent collision energy, which was linearly increased from 20 to 65 eV in positive mode. Low-abundance precursor ions with an intensity above a threshold of 1000 counts but below a target value of 12500 counts were repeatedly scheduled and otherwise dynamically excluded for 0.2 min.

Downstream identification, filtering and validation of LC-MS data were managed using Fragpipe version 24.0. MSFragger version 4.4.0[13] was used for the database search of the LC-MS data versus a FASTA containing all proteins of the *Saccharomyces Cerevisiae* proteome (Uniprot ID: UP000002311) as well as decoy and common contaminant proteins. Search settings were as follows: Trypsin, Enzymatic cleavage at KR, 2 missed cleavages allowed, 20 ppm MS1 and MS2 tolerances, 3 maximum variable mods, Phosphorylation at STY (+79.96633), N-Terminal Acylation (+42.0106), M oxidation (+15.9949). PTMProphet[14] was used for PTM localization and the FDR filter was set to 0.01. Label-Free Quantification was accomplished using IonQuant version 1.11.20[15].

*Spinning disk confocal live-cell imaging*

Cells were grown to mid-log phase in complete synthetic media (CSM) before α-factor (GenScript) was added at a final concentration of 25 µg/mL and cells were incubated at 30°C for 2.5 h. Cells were washed and resuspended in fresh CSM and immediately placed onto CSM agar pads for imaging. Time-lapse imaging was performed on a Nikon Ti-E spinning disk confocal (CSU10; Yokogawa) equipped with a 1.45 NA 100x CFI Plan Apo oil objective, a piezo electric stage (Physik Instrumente), 488 nm and 561 nm lasers, and an EMCCD camera (iXon Ultra 897; Andor Technology). Images were collected using the NIS Elements software (Nikon). The microscope stage was prewarmed to 30°C prior to imaging. Images were acquired every 2 min for 1 h across 21 z-sections collected at 0.5 µm intervals using 100 ms exposure times and 40% laser power for both the 488 nm and 561 nm channels. Z-stacks were maximum-intensity projected in ImageJ and manually analyzed. Myo1-mNG and Htb2-tdTomato were used for analysis. All imaging experiments were performed in three independent biological replicates.

(14) Shteynberg, D. D.; Deutsch, E. W.; Campbell, D. S.; Hoopmann, M. R.; Kusebauch, U.; Lee, D.; Mendoza, L.; Midha, M. K.; Sun, Z.; Whetton, A. D.; Moritz, R. L. PTMProphet:

Fast and Accurate Mass Modification Localization for the Trans-Proteomic Pipeline. *J. Proteome Res.* **2019**, *18* (12), 4262–4272. https://doi.org/10.1021/acs.jproteome.9b00205.
(15) Yu, F.; Haynes, S. E.; Nesvizhskii, A. I. IonQuant Enables Accurate and Sensitive Label-Free Quantification With FDR-Controlled Match-Between-Runs. *Molecular & Cellular Proteomics* **2021**, *20*, 100077. https://doi.org/10.1016/j.mcpro.2021.100077.

### Supplementary Figure 1

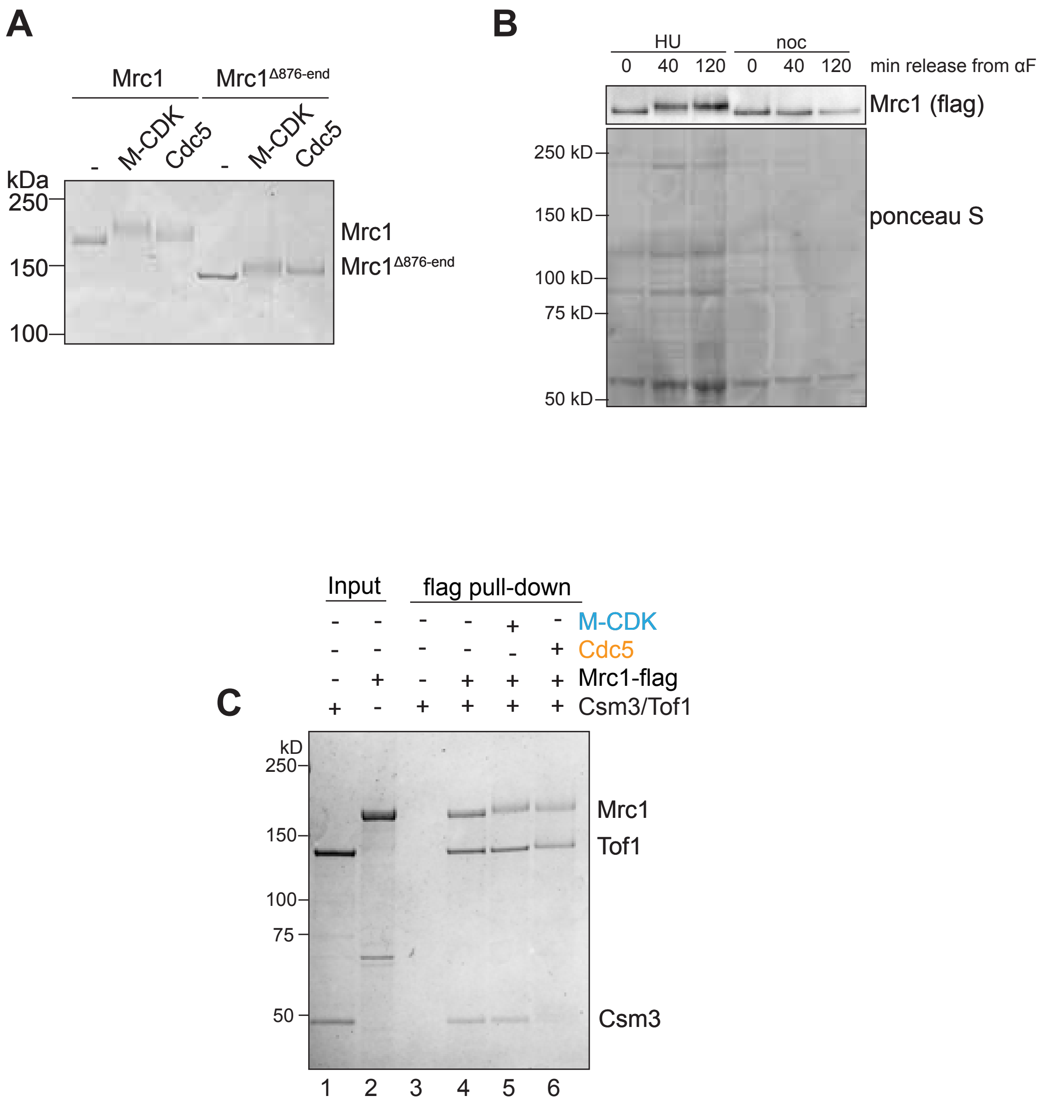


**Suplementary Figure 1.** A) Mrc1 or Mrc1$^{\Delta876\text{-end}}$ were incubated with M-CDK or Cdc5 in the presence of ATP, then run on SDS-PAGE and coomassie stained. B) Cells harboring Mrc1-3xFlag were arrested in G1 with α-factor, then released into fresh media with nocodazole to arrest in prometaphase. Lysates were then immunoblotted with anti-Flag. C) Mrc1 was incubated with M-CDK or Cdc5 prior to addition of Csm3 and Tof1. Mrc1 was then immunoprecipitated with anti-Flag beads and eluted. Samples were then run on SDS-PAGE and coomassie stained.

Supplementary Figure 2

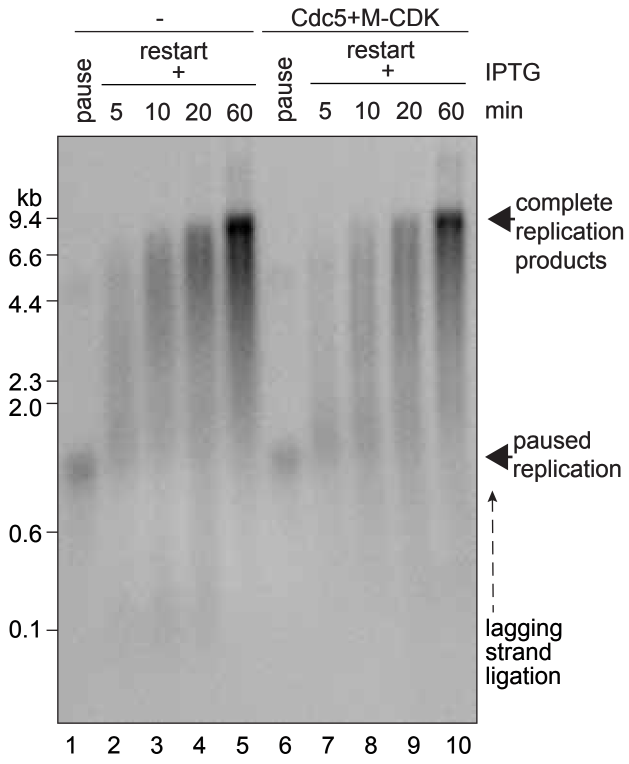


**Supplementary Figure 2.** Experimental setup like Fig 3B. M-CDK were added to LacR blocked replication reactions that also contained Cdc9 and Fen1. Replication was resumed upon addition of IPTG.

Supplementary Figure 3

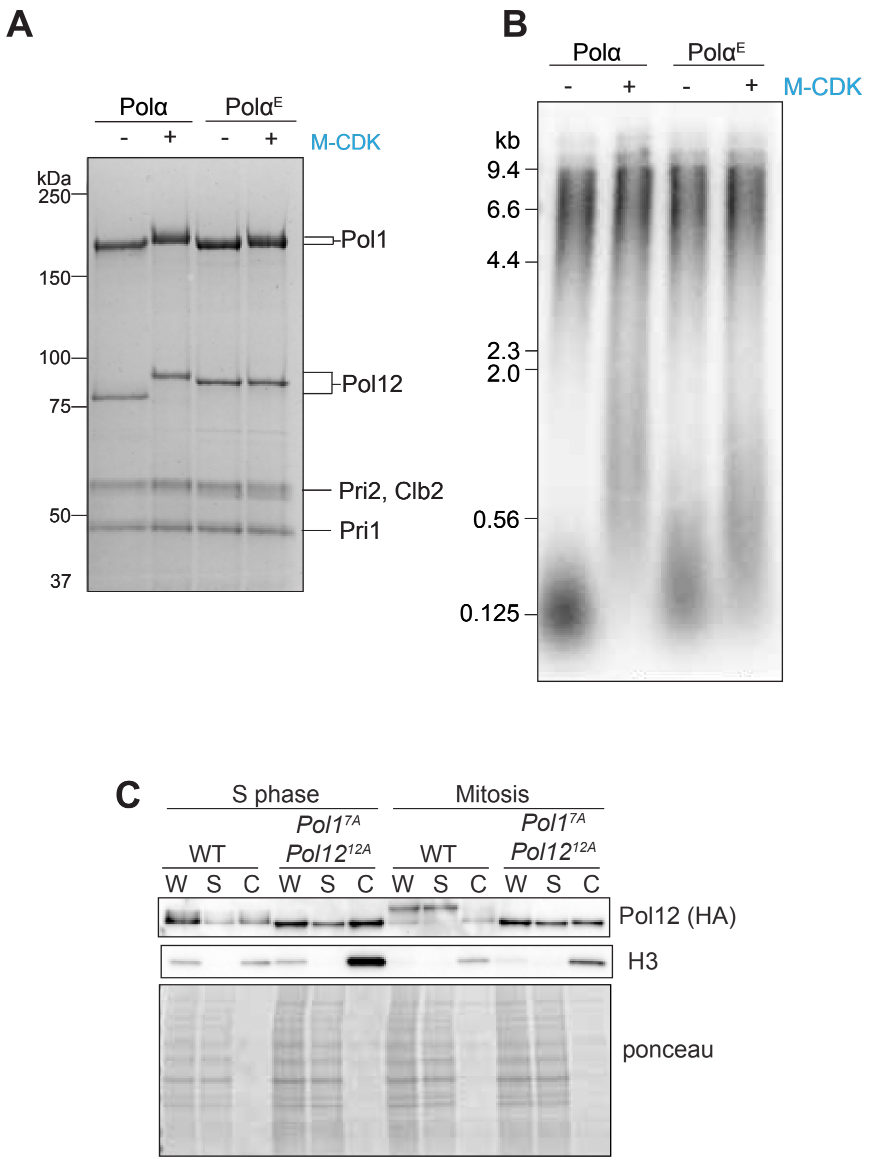


**Supplementary Figure 3.** A) Polα$^{E}$ containing the Pol1$^{7E}$ and Pol12$^{12E}$ mutant subunits was incubated with M-CDK and then separated on SDS-PAGE and coomassie stained. B) Polα$^{E}$ was incubated with M-CDK and then added to a replication reaction. C) Cells from yAWM870 and yAWM902 were arrested in G1 with α-factor for 2.5 h, then released into media containing nocodazole. Samples were harvested 30 min (S phase) and 120 min (mitosis) after α-factor release. Whole cell lysates (W), chromatin pellets (C), and supernatant (S) fractions were processed and then immunoblotted with the indicated antibodies.

Supplementary Figure 4

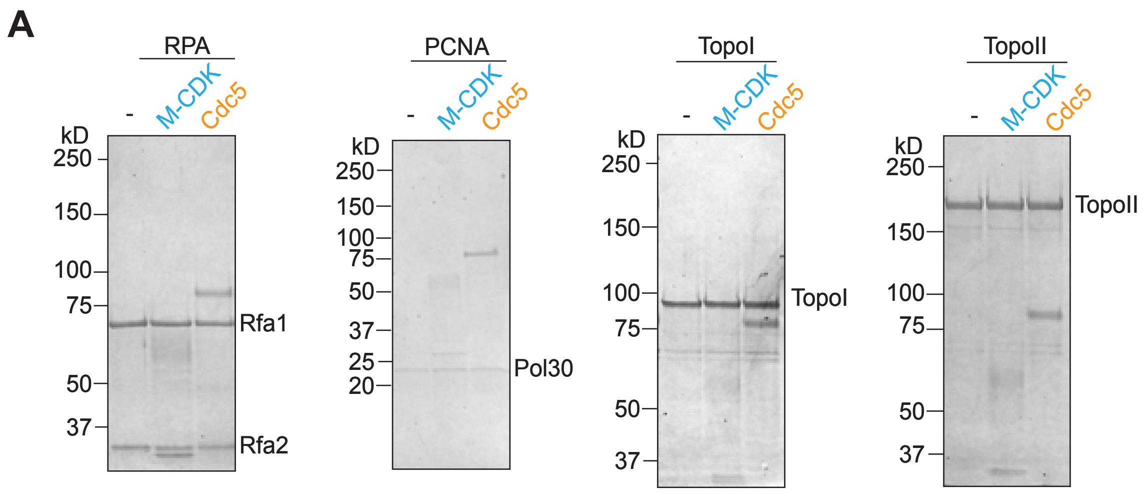


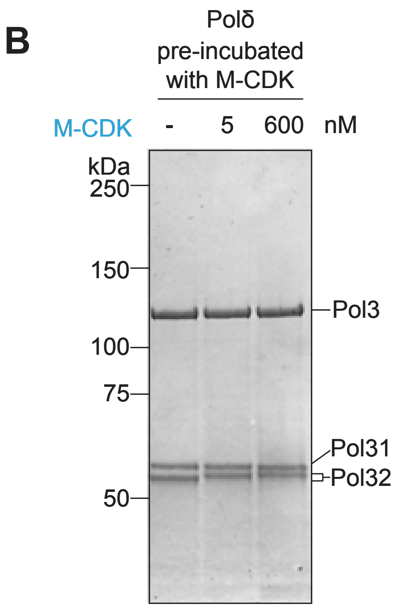


**Supplementary Figure 4.** A) Individual replication fork elongation proteins were incubated 1:1 with either M-CDK or Cdc5 and then separated on SDS-PAGE and coomassie stained. B) Polδ was pre-incubated with 0, 5, or 600 nM M-CDK, then separated on SDS-PAGE and coomassie stained. This Polδ was added to a 20 min replication reaction in Fig 5D.

Supplementary Figure 5

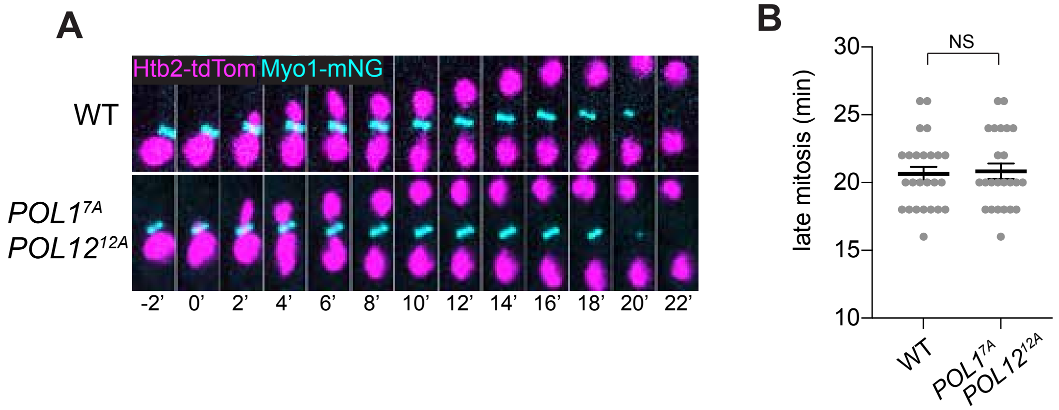


**Supplementary Figure 5.** A) Live-cell confocal imaging was performed every 2 min with the indicated markers. B) Late mitosis timing was determined by nuclear (Htb2-tdTom) entrance into the bud to Myo1-mNG disappearance.
